# Back-illumination Phase Imaging Enables Nanoscale Drift Stabilization in Non-transparent Biological Tissues

**DOI:** 10.64898/2026.03.12.711239

**Authors:** Hanna Manko, Marc Tondusson, Adeline Boyreau, Morgane Meras, Stéphane Bancelin, Laurent Groc, Laurent Cognet

**Affiliations:** Laboratoire Photonique Numérique et Nanosciences, Université de Bordeaux, 33400 Talence, France; LP2N, Institut d’Optique Graduate School, CNRS UMR 5298, 33400 Talence, France; Interdisciplinary Institute for Neuroscience, CNRS, Univ. Bordeaux, 33076 Bordeaux, France

**Keywords:** Microscope stabilization, deep tissue imaging, oblique back-illumination, phase contrast imaging, single-particle tracking, super-resolution microscopy

## Abstract

High-resolution optical microscopy enables nanoscale investigation of molecular structures but is challenged by sample drift during long acquisitions, particularly in thick biological tissues where trans-illumination is unpractical. Precise stabilization at the nanoscale is critical for high-resolution imaging techniques like localization microscopy and single particle tracking. Here we introduce a method combining homogenized differential phase contrast imaging with cross-correlation-based analysis to achieve automated, precise 3D drift correction applicable under oblique back-illumination. We demonstrate its effectiveness in fixed and live organotypic brain slices, maintaining focus within tens of nanometers and enabling high-quality nanoscale mapping of extracellular structures based on single particle tracking. Furthermore, we illustrate its application to opaque liver tissues combined with near-infrared single particle tracking. Our label-free approach provides a versatile solution for stabilizing optical microscopes in thick non-transparent tissues, facilitating extended high-resolution imaging across increasingly complex biological samples.

## Introduction

By enabling studies down to nanoscale in living environments, high-resolution microscopy has become a powerful tool for the investigation of molecular structures and their dynamics in biological tissue ^1–6^. In such optical approaches high-resolution comes however at the cost of long acquisition time. This is the case of localization microscopy techniques which inherently require considerably long acquisition times to gather high number of frames and generate good quality structural data at the nanoscale. For such applications, sample lateral and axial drifts that occur at the nano or microscale can become significant, leading to decreased accuracy of localization and misleading retrieved information. Dynamic studies conducted with nanoscale precisions are also affected by sample movements, as is the case of single particle tracking which is primarily based on the concept of single molecule localization. Therefore, real-time microscope stabilization and/or post-processing drift correction are highly critical especially for super-resolution localization microscopy.

Various methods have been already developed to correct lateral and axial drifts of biologically simple or well controlled samples. The choice typically depends on experimental requirements, such as the sample type, the illumination technique that can be employed and whether drift correction is possible during post-processing. Lateral drifts are the easiest ones to be determined by analysing the lateral movements of sample details in images, which are eventually obtained by ancillary imaging techniques. The latter schematically include label free approaches or require the introduction of contrast agents ^7–10^. On the other hand, correction of axial drifts is more challenging with optical microscopes because images are recorded in orthogonal planes to such movements. A widespread strategy is to monitor the sample axial drift by using infrared light reflected from a coverslip surface. This technique is predominantly used in commercial systems. It is however, only effective for thin samples directly immobilized on a coverslip, such as single layers of cells, that are imaged near an interface displaying notable difference in refractive index to reflect the infrared beam. Another common strategy is the use of fiducial markers, such as fluorescent or scattering nanoparticles, to reveal and correct drifts by analysing their defocusing either in real-time or in post-processing steps ^11,12^. Evidently, fiducial markers must be present in the sample and display photostable signals allowing to track them over the entire acquisition time. In thick samples, introduction of such objects in any arbitrary plane can turn out to be extremely difficult, if not impossible.

Some efficacy in automated refocusing was demonstrated in transparent samples through the use of cross-correlation (CC) of bright-field images of the unlabelled sample to monitor the sample drift ^7,13^. Differential phase contrast (DPC) imaging was shown to provide enhanced structural contrast and improved optical sectioning ^14^. However, conventional implementations rely on trans-illumination and are therefore limited in thick or highly scattering tissues ^15^. Indeed, their densely packed and highly overlapping structures, result in low level of details in bright-field imaging, and image contrast quickly vanishes due to tissue absorption and scattering making this approach inappropriate. A strategy which can effectively detect small axial drifts in thick tissues using back-illumination would be highly desirable to ensure high performance across a wide range of sample types.

Here, we introduce homogenized differential phase contrast (hDPC) oblique back-illumination imaging, which enables detailed label-free visualization of opaque deep tissue structures in a back-illumination configuration. We further show that this approach generates unique performance for cross-correlation-based autofocusing. When combined with standard methods for correcting lateral drifts, we demonstrate that this approach can effectively compensate for 3D micro- to nanoscale sample drifts in single-particle tracking experiments performed in both fixed and live brain tissues, using not only back-illumination but also trans-illumination when tissue transparency allows for it. Finally, we show its effectiveness in thick liver tissues, where obtaining label-free images is feasible only under oblique back-illumination.

## Results and discussion

Label-free imaging techniques are essential for long-term observation of biological samples such as cells and tissues. However, generating high-contrast images of intrinsic tissue structures typically requires the use of methods allowing contrast enhancement such as phase contrast or differential phase contrast microscopy ^14^. Oblique illumination techniques provide lateral phase-gradient contrast, which produces images resembling those of DIC microscopy and can reveal tissue structures in greater detail ^16^. However, all these techniques rely on trans-illumination modality which limits their applications to thin, nearly transparent samples. Oblique back-illumination (OBM) was introduced to enable label-free imaging inside thick tissue ^17^ allowing thick sample illumination from the same side as detection is performed by using non-coherent illumination source like light emitting diodes (LEDs). This configuration enables multiple scattering in thick tissue, making it possible to mimic trans-illumination at the focal point by detecting back-scattered photons leading to phase-gradient contrast.

By sequentially illuminating the sample with opposing light sources offset from the detection axis and subtracting the two resulting images, one can enhance the phase gradient, suppress absorption contrast, and achieve tomographic sectioning. These differential phase contrast (DPC) images provide a relief-like appearance of tissue structures while maintaining strong directionality. Further development of this technique exploits illumination by four independent sources leading DPC images along two orthogonal directions (Figure 1a). Quantitative OBM (qOBM) enables phase reconstruction trough a Tikhonov regularized decon-volution of two orthogonal DPC images, which requires the knowledge of transfer function of the system ^18,19^. This step is computationally intensive, and because the transfer function depends strongly on tissue-specific scattering properties, it may not be easily generalized between samples, which limits the use of qOBM in autofocus applications. In Figure 1, we present two phase-contrast images obtained inside fixed organotypic brain slice using two opposite light sources and two additional sources positioned orthogonally (Figure 1b,c). It is evident that the resulting images display pronounced directionality in the relief structures, which can vary considerably depending on the region in the sample (Figure 1c).

**Figure 1.**
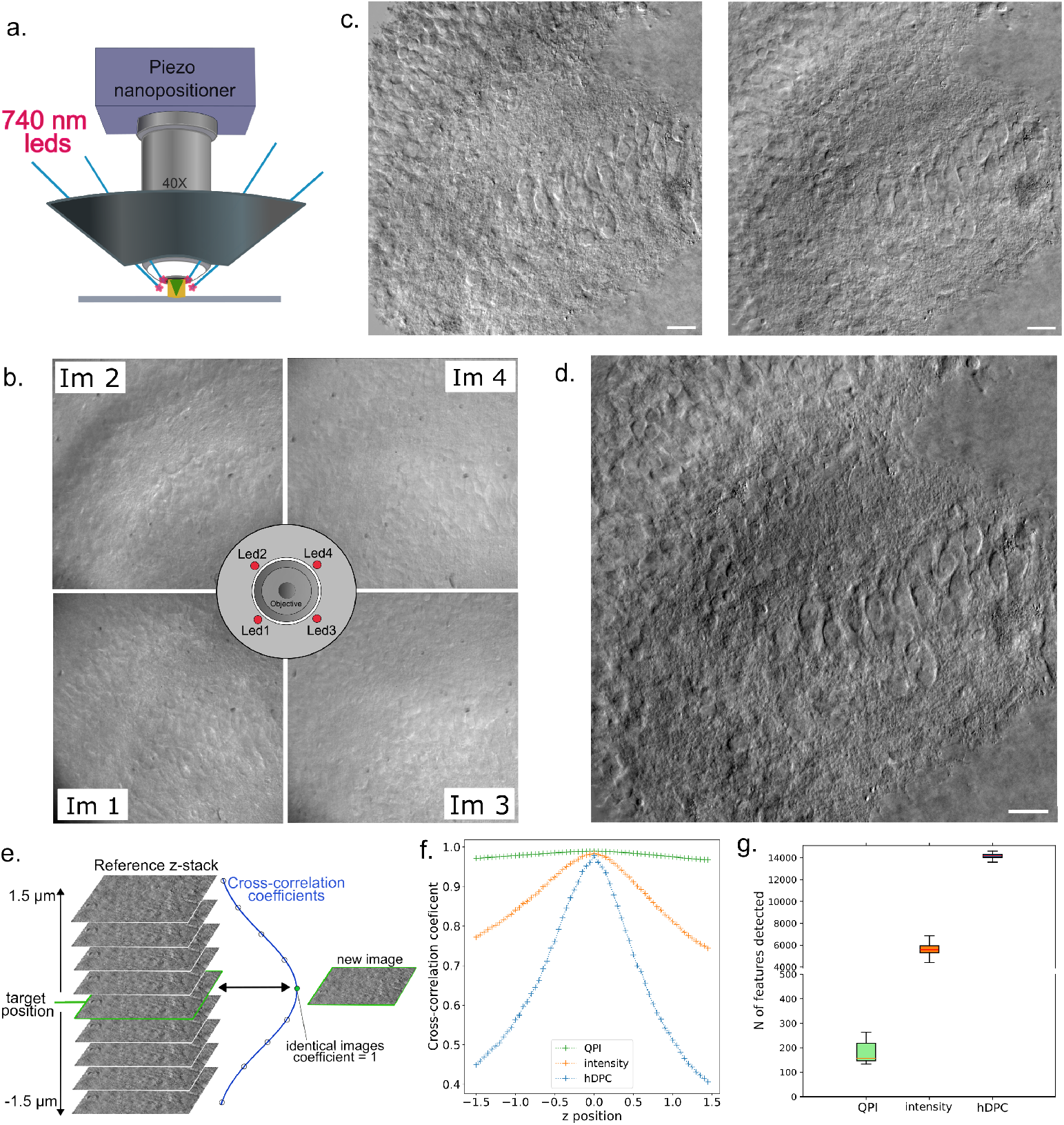
**a.** Schematic of the fiber positions around the microscope objective, held in place using a custom 3D-printed holder. **b**. The 4 raw images obtained while illuminating the sample with 4 fibered LEDs positioned as shown in the middle. These images are used to obtain two DPC images **c**. and finally the hDPC image **d. e**. The z-stack of images acquired at different focus planes around the target z=0 position in the range -1.5*µ*m : 1.5*µ*m. The CC coefficient reaches its highest value for the middle image and gradually decreases in both positive and negative directions when calculated between the middle image and each image in the z-stack. **f**. The cross-correlation curves obtained by calculating normalized CC coefficients between image at z=0 and every image in z-stack acquired for intensity, phase, and gradient images (Figure S1) obtained using trans-illumination setup (Figure S2,a). **g**. The number of features detected on the different types of images. The values were obtained for z-stacks acquired in three different fields of view.

**Figure 2.**
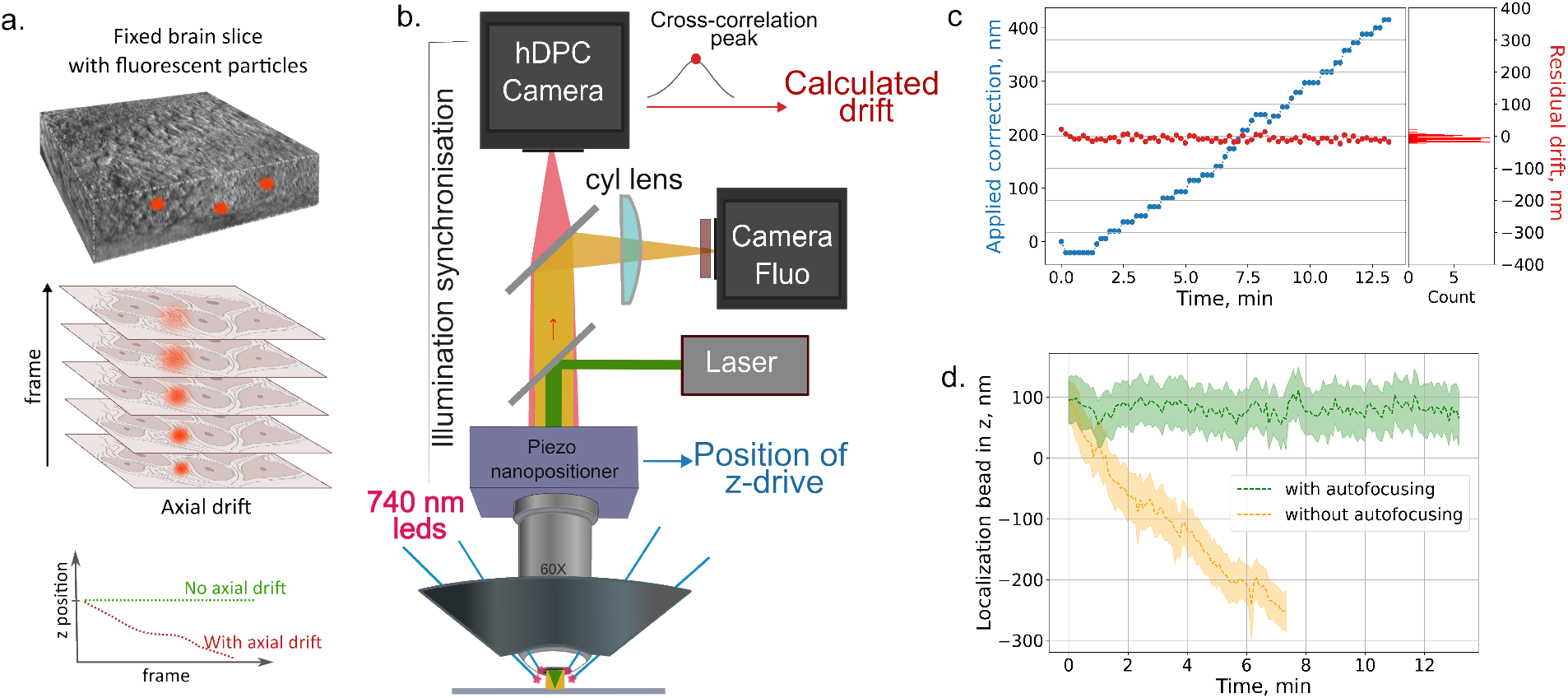
**a.** Immobilized fluorescent particle in a fixed brain slice and the expected behavior in the presence of focus drift during acquisitions.**b**. Schematic of optical setup using oblique-back illumination. **c**. During autofocusing the drift was calculated for every 5^*th*^ frame acquired using cross-correlation of gradients images (red curve, axial drift obtained right before its correction). Additionally, the current z-drive position of the microscope was recorded (blue curve). **d**. The bead axial position recorded with and without active stabilization. The standard deviation was calculated using sliding window of 100 frames.

To eliminate illumination-direction dependent variations in image detail, we propose here to combine the DPC images acquired along both the x- and y-axes. This resulted in a homogenized DPC (hDPC) image in which structural details are independent of the illumination geometry or sample orientation and which contain high amount of structural details (Figure 1d). In the case of OBM, reference images (Im_*Ref*_ ) acquired without sample, are required to compensate for variations in the illumination provided by each individual LED for each image (Im_*Raw*_). Thus, to obtain hDPC we use:

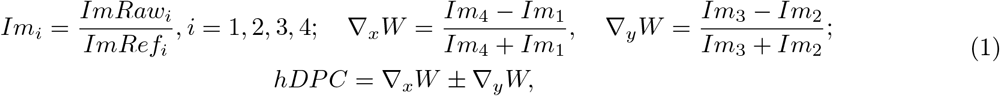

where *i* - image index which corresponds to LED number as displayed on Figure 1d. The ± operator indicates plus or minus, which may depend on specific setup and some variations in illumination provided by LEDs.

We postulate that such highly detailed images with improved sectioning capabilities would be particularly beneficial for automated refocusing in thick tissues, as the method relies on the intrinsic structure of the sample. We thus first assessed this method by recording a set of images recorded at a series of precisely defined focus levels around the target plane position that act as reference points (reference z-stack). Further, during an acquisition, each new image is compared to images in the reference z-stack by calculating normalized CC coefficients. The highest correlation is obtained for the most similar image in z-stack and the target focus is thus at the maximum of the CC curve (Figure 1f). Since images are recorded at well-defined z-planes, the current axial position can be determined by the location of the correlation peak while direction is directly encoded in axial coordinates of CCC peak.

We applied this procedure to a stack of intensity images obtained inside a ∼ 100 *µ*m thick brain slice in the range of 3 *µ*m around the focal plane with 50 nm step, and for each image in the reference z-stack we calculated the CC coefficients against the middle image. The width of the CC curve and consequently the effectiveness of the method is highly dependent on the amount of features in the image, and their variation with defocus, both being dependent on the NA of the objective used but also on the imaged sample which can display specific absorption and scattering characteristics thus affecting the image contrast variations. In the field of imaging, features are typically defined as identifiable patterns, intensity variations or structural elements that serve as distinctive landmarks within an image ^20^. Image normalization reduces the dependence of the calculated CC coefficients on illumination variations by removing global intensity offsets and scaling effects. As a result, the CC coefficient will be more sensitive to pixels for which *I* ≠ *I* (I - pixel intensities), implying that pixels carrying structural information (specific “features”) contribute more strongly than homogeneous, textureless regions. Consequently, images containing a higher density of features or fine details produce sharper CC peak. Furthermore, the peak of the cross-correlation function is fitted using a polynomial model allowing for subpixel determination of the peak position. Because the peak position is derived from a polynomial fit to the correlation curve, it represents a weighted estimate that depends on the correlation coefficients in the local neighborhood of the maximum.

Organotypic brain slices are standard *in vitro* imaging system for their relative simplicity to handle as compared to *in-vivo* organisms ^21^ making them widely used for imaging or electrophysiology studies for instance. The choice to perform the first demo on organotypic slices allowed us to benchmark hDPC with regular bright-field imaging techniques. Such methods as quantitative light shift interferometry (QLSI) allows to obtain two directional DPC images that can be generated prior to phase extraction (see Supporting Information, section 2).

This enables comparison of CC-based method performance applied to different image types, such as intensity-based, hDPC or QPI images (see Figure S5), when applying the cross-correlation (CC)-based method. We find that hDPC displays significantly more details than QPI and also than intensity images (Figure 1g). Furthermore, the improved optical sectioning capabilities of hDPC images are evident in the higher number of matching features detected across defocus levels (see Supporting Information, Figure S3). Accordingly, the highly pronounced CC peak obtained from hDPC enables an efficient discrimination between correlation coefficients, even for images with minor changes in focus (Figure 1f, blue curve) which is a clear asset for precisely correcting axial drifts. Moreover, higher number of detectable features on hDPC results in better xy-drift correction using methods such as scale-invariant feature transform (SIFT) which was used in this study.

### Stabilization in fixed brain slices

In order to test the performance of hDPC under oblique back-illumination to correct 3D sample drifts in real time, we first applied the approach to an experiments of 3D single particle tracking (SPT) performed in epi-configuration inside fixed organotypic mouse brain slice where 100 nm fluorescent nanobeads were introduced (Figure 2,a). Fixed brain slices broadly serve as relevant imaging models as they preserved structural organization of the original brain tissue. In this samples, all structures are static and some immobilized fluorescent beads could be identified at the depth of 5-15 *µ*m from the slice surface. Even with a static sample, drift may occur from factors like temperature fluctuations, vibrations, or mechanical relaxation. To achieve subwavelength super-localization of immobile nanobeads, we implemented the standard astigmatism method by inserting a cylindrical lens into the light collection path (Figure 2,b). This lens induces a z-dependent distortion in the point spread function (PSF), allowing axial super-localization of the nanobeads thrpugh analysis of the PSF’s ellipticity and orientation ^22^. Monitoring the axial position variations of the immobile nanobeads thus allows validating the effectiveness of our autofocus method based on hDPC images.

**Figure 3.**
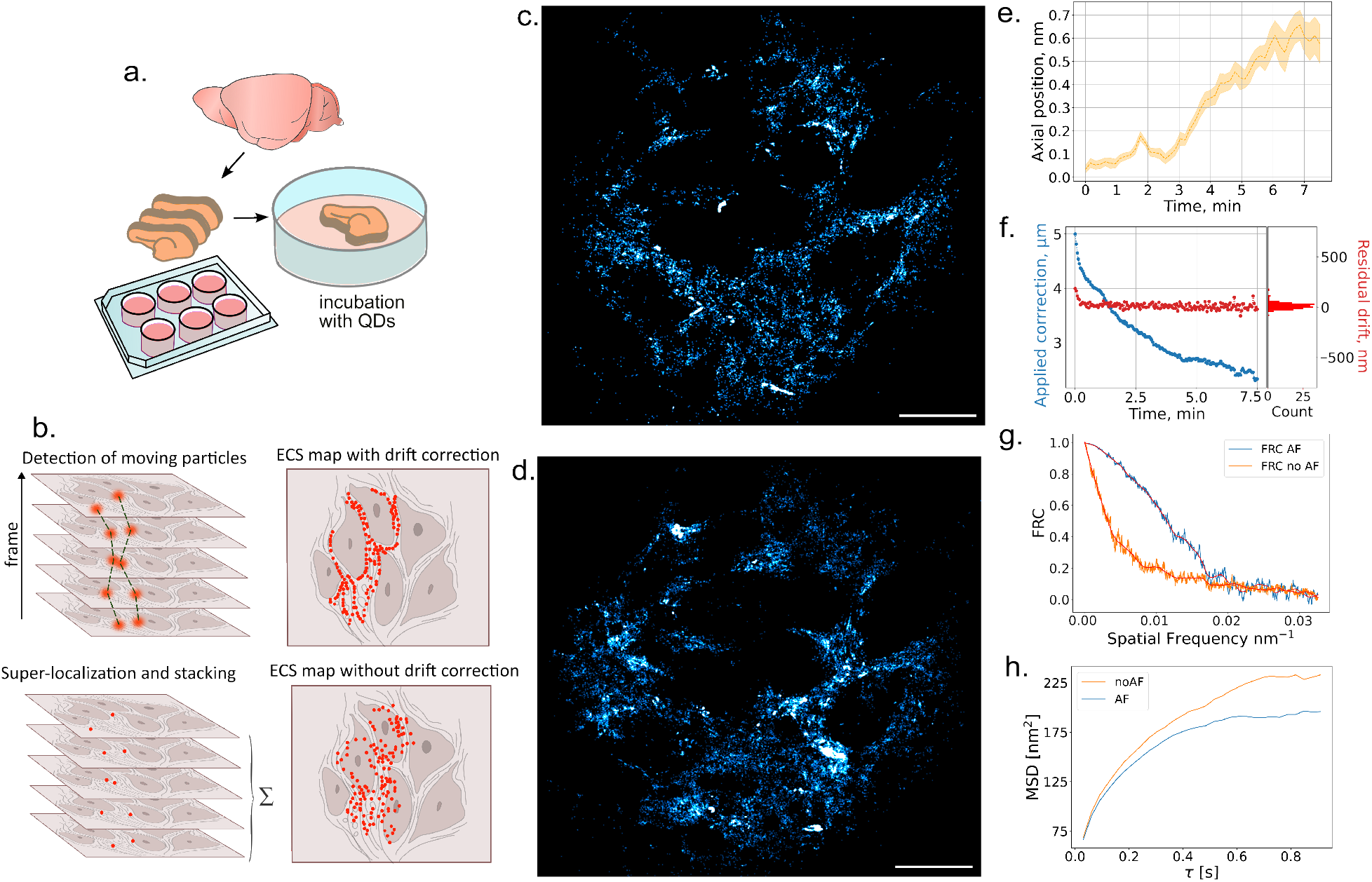
**a.** Organotypic brain slices preparation. **b**. Generation of ECS maps. The reconstructed image of QDs localizations in ECS at ∼ 25 *µ*m inside brain tissue without active autofocusing system **c**. and with **d**. (scale bars = 10 *µ*m) for back-illumination setup. **e**. The drift recalculated at the post processing step using the hDPC recorded simultaneously with QDs tracking without active autofocusing was ∼ 600 nm. **f**. The drift detected (red dots) with SD value of ∼ 26 nm and the position read from z-drive (blue dots) using hDPC obtained via back-illumination while total corrected drift was 2.5*µ*m -blue dots. **g**. The FRC curves calculated for the images obtained with active autofocusing and without. **h**. Mean-squared displacements of QDs with and without active autofocusing.

In practice, the procedure of the 3D stabilization of the microscope contains several steps. For every selected region of interest (ROI) and focus position, a reference z-stack around the target position needs to be acquired only once. Typically, the reference z-stack is acquired in the range of -1.5:1.5 µm with 50 nm steps. Typical time required for acquisition of such z-stack was 12 s. Once the acquisition with autofocusing is started, the xy-drift need first to be corrected for every n^*th*^ acquired image (depending on the chosen autofocusing frequency). For this purpose, the scale invariant feature transform (SIFT) ^23^ method is applied to ROI of 700 x 700 pixels and lateral drift is corrected numerically. The numerical correction applied before each autofocusing step helps prevent backlash errors caused by the xy-motion of the stage. Therefore, mechanical correction is only necessary in cases of significant xy-drifts (more than 50 µm). Using SIFT-based lateral drift correction on DPC images, it is possible to correct xy-drift in the corresponding fluorescent channel with a precision of ∼ 14 nm (Supporting Information, section 4, Figure S4). Further, the normalized cross-correlation coefficients are calculated between ROIs of every n^*th*^ image and images in reference z-stack. The obtained cross-correlation curve is fitted using polynomial fit (whose degree can be adjusted in real time). The drift ( Δz) is calculated as difference between target position (arbitrary chosen to be at z=0) and the current position, obtained by determining the peak position of the fit. The axial drift is then corrected by acting on the z-drive of the microscope (Figure S5). A practical approach for performing acquisitions with such an autofocusing system is to display the CCC curve obtained at each autofocusing step, allowing real-time monitoring of changes in the curve shape. This strategy was implemented in our Python-based software (see Materials), enabling visual inspection of the generated curves. When distortion becomes significant - for example, when the peak becomes highly asymmetric or noisy, it is necessary to update reference z-stack (see also Supporting Information, section 5).

The hDPC were obtained while illuminating the sample with 740 nm fiber-coupled LEDs, placed around the microscope objective at 45° from the sample surface. This wavelength was chosen to minimize the light absorption by the tissue, while producing good scattering properties as required in OBM ^24^. During the test we were able to maintain the focus position in range of ±15nm (Figure 2c, red dots), with a standard deviation of the measured drift of 9 nm. The calculated SD of the mean axial z-position of the bead was determined to be 10 nm (Figure 2,d). The minimal axial drift corrected by the piezo was intentionally limited to 10 nm to avoid overshooting and maintain stability. As a result, the stepping of the position recorded from the z-drive can also be observed (Figure 2c, blue dots).

In such samples, similar performance was achieved in trans-illumination configuration by applying CC-based method to hDPC images derived from QLSI method on an inverted microscope (Supporting Information, section 2, Figure S2).

### Stabilization in live slices

#### Organotypic brain slices

We next aimed to test the performance of the autofocusing system in live samples. For this purpose SPT of CdSe/ZnS biocompatible quantum dots (QDs) was performed in organotypic rat brain slices (7 days *in vitro*), maintained at 35°C on the microscope (Figure 3a). The incubation of the slice with QDs allows them to efficiently navigate in the extracellular space (ECS) and by localizing their positions a super-resolved map of the ECS architecture can be generated ^25^ (Figure 3b). For this, a high number of localizations is necessary, and thus long acquisition times are needed.

In the following, QDs were identified and tracked at a depth of around 25-50 *µ*m inside the tissue slice. The ROI is typically chosen using hDPC images and further adjusted using the signal from QDs in the fluorescent channel to capture an area with multiple nanoparticles diffusing. A common way to minimize the impact of the drift in SPT experiments consists in limiting the number of consecutive frames acquired while tracking the nanoparticles in the brain ECS ^25,26^. While increasing the numbers of acquired frames to several thousands the axial drift can indeed become significant.

Here we acquired ∼ 15000 frames with exposure time of 50 ms to generate a high number of QDs localizations to reconstruct ECS maps (Figure 3c,d). Figure 3c, shows that a defocus of only 600 nm (Figure 3e) is sufficient to affect the super-resolved maps generated by QD localizations in the brain ECS. In contrast, Figure 3d and f, shows that active autofocusing enable maintaining the focus position when overall axial drift is 2.5 *µ*m which in some cases can reach up to ∼ 10 *µ*m (Figure S7). Not surprisingly, axial drift SD values in live sample were found to be higher than those for fixed tissues. This difference can likely be caused by more pronounced focus drift due to greater impact of maintaining the slices at physiological temperature and continuous changes within the tissue.

Beyond the qualitative description of the improved quality of ECS maps obtained with autofocusing, we quantified the impact of sample drifts on the quality of the ECS structures revealed by the QD super-localization maps. The task is difficult as QDs are continuously moving and sample drifts are intricated into the QDs. While we do not anticipate change in the particle localizations, we postulate that in the ECS maps, which consist of the assembly of thousands of QDs superlocalizations, focus drifts introduce some blurring of the ECS structures if QD localizations coming from different focal planes are pulled together.

To test this hypothesis, we applied Fourier Ring Correlation (FRC) which is commonly used to quantify the resolution of super-resolved microscopy images, mainly for well-defined static structures such as microtubules in fixed cells ^27^. In our case, since the ECS maps are extracted from particles trajectories, the FRC curve will be strongly affected by the drift that result in heterogeneous images containing structures from several planes. As expected, the resolution is similar with and without autofocusing. In contrast, the FRC values at medium and high spatial frequencies are strongly decreased without autofocusing, highlighting the blurring effect of the drift. Hence, the FRC analysis applied to our data, and notably the change in area under the curve, indicates an increased quality of the ECS maps obtained with autofocusing (Figure 3g). Finally, we calculated the mean-squared displacement (MSD) of the QD trajectories to evaluate how focus drift influenced the measurements. We found that a drift as small as 600 nm is sufficient to distort the MSD curve, leading to an overestimation of the diffusion rates and explored areas (Figure 3h). Note that despite their thickness, brain slices are still sufficiently transparent to allow experiments with a trans-illumination setup allowing us to observe similar performance of the automatic refocusing procedure (Figure S8).

#### Liver slices

We finally aimed to demonstrate the effectiveness of the proposed method on opaque samples like liver slices. Liver slices are interesting ex vivo model, as they preserve the structure and microenvironment of the whole livers. As such they are proven to be suitable for disease modelling, for example liver fibrosis, which interstitial space has been shown to be altered in the early stage of fibrosis propagation based on the analysis of the diffusion properties of nanoparticles emitting in the short wave infrared (SWIR) ^28^.

Due to its intrinsic opacity and high blood content, the liver exhibits pronounced scattering and autofluorescence. These optical properties prevent reliable SPT of visible emitters, including quantum dots. Therefore, for SPT experiments in liver tissue, we have previously introduced the use of single wall carbon nanotubes (SWCNTs) as fluorescent probes. Their high brightness and photostability, combined with the ability of specific chiralities to be excited and detected in the infrared, enable deep-tissue imaging even in strongly scattering organs like the liver ^28^. Moreover recent development of ultrashort Color-Center bearing NanoTubes (uCCNTs), which combine bright and photostable SWIR emission with reduced dimensions, showed facilitated diffusion in the brain’s extracellular space^29^, making them the perfect candidate for experiments in opaque livers.

In present work we used fresh liver injected with uCCNTs and sliced right before the imaging (Figure 3a). Because the typical size of such slices is much larger than brain slices, approximately 2.5 cm length and 600-800 *µ*m thick the stable immobilization of the slice on the glass coverslip surface if more challenging. The addition of immersion liquid on top can further induce unexpected movements of the slice during acquisition. Consequently, sample drifts are not slow nor primarily directional, as observed in brain slices, and, in some cases, characterized by sudden, unpredictable jumps (Figure 4 blue dots).

**Figure 4.**
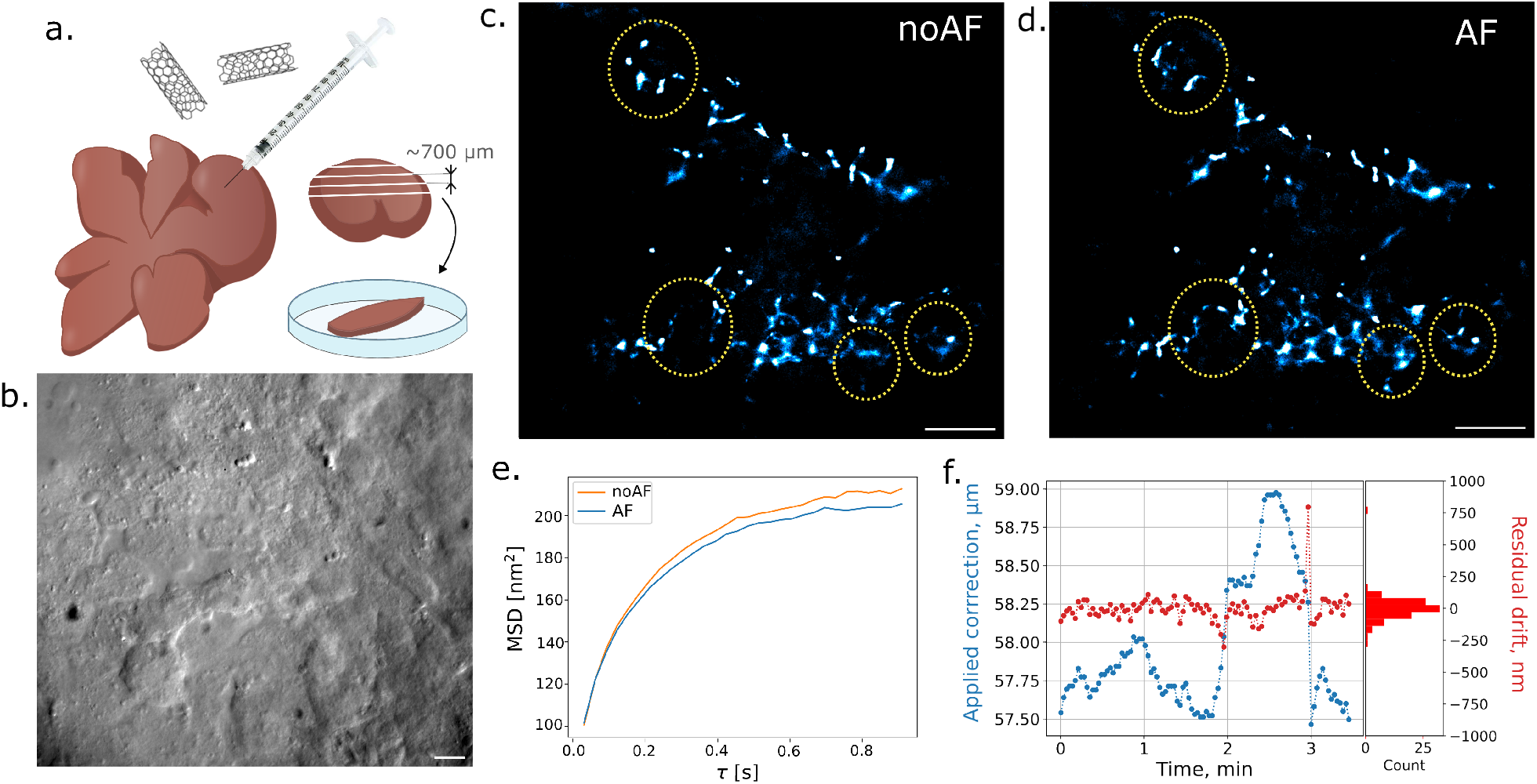
**a.** Liver slices preparation. **b**. hDPC image of live liver slice tissue (scale bar = 20 *µ*m). The liver slice ESC maps obtained without **c**. and with **d**. active autofocusing (scale bars = 10 *µ*m). The differences between the ECS maps are clearly visible, with several regions highlighted by circles. The area marked in yellow contains finer structural details in the ECS map reconstructed from the acquisition with active autofocusing. In contrast, the region indicated in green shows an area where the nanoparticles explored a different portion of the extracellular space. **e**. The corresponding MSD curves. **f**. An example of drift curves obtained in live liver slices, showing that the drift is non-directional and exhibits strong variations over time.

The acquisition time in liver tissue was limited to approximately 5 minutes, as morphological changes occur faster than in brain slices. A first observation from the reconstructed maps of nanoparticle exploration with active drift correction is that they contain richer structural information, revealing features that are not visible without drift correction (Figure 3c,d). We next attempted to perform FRC analysis on the ECS maps from liver slices; however, this was not feasible because the structures present are more discontinuous than in brain slices making this analysis inappropriate (Figure 3c,d). On the other hand, the calculation of MSD curves reveals that sample drift leads to an apparent overestimation of the diffusion rates and explored areas of SWIR uCCNTs, similar to what was previously observed for QD diffusion in brain slices.

## Discussion

We have shown that application of the cross-correlation-based method to hDPC images acquired under oblique back-illumination is an effective approach to perform automated refocusing in thick tissues. The combination of autofocusing system with existing methods for xy-drift determination allows to stabilize the microscope in 3D with precision down to 14 nm. The use of hDPC images generates well-defined CC peak which provides high accuracy of the determination of the current axial position. This is ensured by the higher detected features and stronger variation of image content in comparison to intensity only or QPI images. The application of CC-based method to hDPC then allows to effectively detect and correct the focus drift in both fixed and live samples in real time. Furthermore, the drift correction precision could be improved during post-processing stage while using the same hDPC images.

As hDPC images can be generated using a variety of label-free techniques, it is possible to apply the method to a wide range of sample types and to use different illumination modalities, including oblique back-illumination, making it now possible to stabilise optical microscope in thick, non-transparent tissue with nanoscale precisions which was not possible with existing techniques. Moreover, the method can be adapted to various illumination wavelengths, enabling its use with a broad variety of fluorophores ^30^. The method developed in this study is specifically designed to correct global sample drift, as our particle tracking experiments aim to extract detailed diffusion properties of particles within static, localized regions of thick tissue. For experiments requiring information on longer trajectories or particle motion across larger tissue volumes, additional approaches, such as azimuthal tracking ^31,32^ or extended-focus methods ^33^ - are required. However, even in these cases, correcting for coordinate offsets caused by sample drift or tissue motion relative to particle movement remains essential.

In this work we typically limited acquisitions to around 10 min for brain slices and 5 min for liver slices. Indeed, over longer time intervals the effectiveness of the proposed autofocusing method (as well as the relevance of performing SPT and generating ECS maps) might be constrained by the morphological changes in live tissue, which will alter the shape of the CC curves and therefore destabilise the autofocusing process. In such situation, it is required to record new reference z-stack and restart the acquisition. Nevertheless, the application of CC to hDPC enables the acquisition of much longer movies even in such conditions, while ensuring the imaging at the precise focal plane in thick tissues, exceeding previous capabilities

On a broad scope, we foresee that our method can be deployed in a large number of microscopy configurations for a variety of applications such as in biological or material science.

## Methods

### Sample preparation

#### Fixed organotypic brain slices

Hippocampal organotypic slices from postnatal 5-days-old Sprague-Dawley rat pups, both male and female (Janvier/Charles River, France) were prepared as previously described ^25,34^. The slices were fixed using 4% solution of paraformaldehyde in PBS, containing 1:1000 dilution of Tetraspec beads (4-color, 0.1 *µ*m, T7279 - Thermofisher Scientific).

#### Live organotypic brain slices

For SPT experiments the commercial photoluminescent Qdot™ 655 (Q11422 MP, Invitrogen) has been used. The slices were first washed with HEPES-based aCSF for 10 min to remove culture medium and then incubated with QDs suspension (120 pM) in HEPES-based aCSF solution containing 1% BSA for 40 min at 35°C. After washing for 10 min the samples were imaged with the sample temperature controlled at 35°C (Tokai Hit).

#### Live liver slices

The liver slices were prepared as previously described ^28^ The solution of uCCNT was injected into left lobe with 0.6×0.25mm sterile needle (Agani, Terumo). The lobe was then embedded in low-melting point agarose (Thermo Scientific) gel and cut into 650 *µ*m slices with vibratome (Leica VT1000S). The slices then were places into imaging chamber with thin layer of agarose gel and then imaged in PBS with the sample temperature controlled at 35°C (Tokai Hit).

### Oblique back-illumination optical setup

A custom-built modular microscope (Thorlabs) was used for oblique back-illumination. Four 740 nm LEDs were injected in fibers (Thorlabs, FP1000ERT) and mounted around the microscope 60X 1.0 NA objective (Nikon) using 3D printed fiber holder. The fluorescence emission was separated from LED light by 705 nm dichroic mirror (FF705-FDI01, Semrock). The detection in fluorescent channel was performed by Hamamatsu (ORCA-Flash 4.0) camera and controlled separately by Micro-Manager software ^35^. The LEDs illumination was synchronised with the second camera (KURO®, Princeton Instruments) exposure using hardware triggering.

The control of the LEDs was performed using Arduino Uno board. The Arduino board, hDPC channel camera and piezo z-nanopositioner (FOC.100, Piezoconcept) were controlled by custom Python software with dedicated user interface. The same software was used to perform the acquisitions required for autofocusing algorithm. The procedure for connecting all the elements is described in the user manual, which is available along with the dedicated software.

For imaging Tetraspec beads the 638 nm diode laser was used together with a dichroic mirror (Di01-R635, Semrock) and a band-pass filter (BP 685/40, Semrock) placed before fluorescent channel camera. A cylindrical lens (f=1000 mm) was introduced into the fluorescence detection path to extract the z-coordinates of the particles. A cylindrical lens calibration was obtained right before the acquisition using the same bead. The QDs were imaged using 488 nm laser for excitation and a dichroic mirror (Di02-R561, Semrock). A 655/15 single-band bandpass filter (BrightLine®) was used to filter emission in fluorescent channel and a 500 nm longpass filter which was placed before the QLSI module.

For liver experiments we used 40X 0.8 NA objective (Nikon). The LED light was separated using long-path 900 (DMLP900R, Thorlabs) dichroic mirror while using 721/65 (FF01-721/65, Semrock) filter for additional filtering. 985 nm (AeroDiode) laser was used for excitation of uCCNTs. The light was reflected from dichroic mirror 1064 nm (LPD01-1064RS, Semrock) and the emission from the sample was further filtered by 1100 nm filter (FELH1100, Thorlabs) before beeing detected by C-RED 2 (First Light imaging) camera.

### Lateral drift correction

In this study, xy-drift was corrected at each autofocusing step. To prevent additional movement of the slice in the imaging medium and avoid backlash errors from stage adjustments, the correction was performed numerically. The drift correction utilized the SIFT (Scale-Invariant Feature Transform) method available in the silx.sift Python library. This approach allows for lateral drift correction at the subpixel level, particularly when the image contains a high amount of features. For more details, refer to the documentation: https://www.silx.org/doc/silx/latest/Tutorials/Sift/sift.html.

### Autofocusing procedure

For the back-illumination the reference is built by averaging a set of images in z-range of -45:45 *µ*m with 10 *µ*m steps.

Each image in the reference z-stack has a well-defined axial position, as the reference z-stack is acquired at precisely controlled z-positions, typically spanning -1.5:1.5 *µ*m with a step size of 50 nm. Thus, if an axial drift of the sample occurs during acquisition (for example -200nm), the newly acquired image at this displaced position will be more similar to the reference image in z-stack recorded at the corresponding position. As a result, the CCC reaches its maximum at around z = -200nm. The axial coordinates of this CCC peak, retrieved upon polynomial fitting, therefore provides both the magnitude and the direction of the actual drift. The normalized CCC are calculated as :

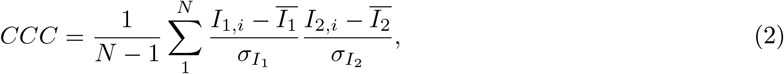

where I – pixel values of the image, N – image size, *σ* -standard deviation. In order to increase the speed of the reference stack acquisition, one can use uneven stepping of the z-drive during z-stack acquisition with the intervals smaller (50 nm) around the desired (middle) position and larger further from it.

The calculations required for each autofocusing step were performed using a smaller ROI, typically 700×700 pixels. During the experiments, GPU acceleration was used only for the XY-correction step of the SIFT method. Microscope control and all calculations were carried out on a computer with 64 GB of RAM, an AMD Ryzen 7 7700X processor, and an NVIDIA GeForce RTX 4060 GPU. The average time required to perform the calculations for each autofocusing step was approximately 0.7 s. However, a deliberate time delay was introduced during the experiments, resulting in ∼ 1.5-2 s intervals between each autofocusing step (see Supporting Information, section 5.1). The calculation speed can be further increased by using an even smaller ROI, performing all calculations on the GPU, and implementing parallelization for certain steps.

### Data analysis

The localizations of fluorescent particles were retrieved using ThunderSTORM ^36^ plugin in Fiji and further analysed using custom Python code.

For the FRC analysis, a similar number of localizations was used for both conditions, with and without autofocusing. Each data set was split into two subsets, one containing the localizations with odd indices and the other with even indices from the ThunderSTORM results table. The FRC curves were further obtained for reconstructed images based on Normalized Gaussian using Fourier Ring Correlation Plugin in Fiji.

## Acknowledgements

The authors thank S. Nandi for experimental support, R. Mangalwedhekar for discussions and P. Teulat for mechanical parts fabricated for this project.

## Data and Software Availability

Data underlying the results presented in this paper are not publicly available at this time but may be obtained from the authors upon reasonable request.

The custom python-based software and the user interface developed during this study together with user manual are freely available at https://github.com/hmanko/hDPC_autofocus.git.

## Supporting information

The following files are available free of charge.

- Section 1: Comparison of real-time autofocusing methods; Section 2: Autofocusing using trans-illumination, Figure S1: An example of intensity, phase and hDPC images obtained using QLSI module and Figure S2: Schematic of optical setup for trans-illumination and results of autofocusing performed in fixed slices; Section 3: Features sensitivity to defocus and Figure S3 - Number of matching features Section 4: xy-drift correction and Figure S4 - comparison of xy-drift correction using fiducials and hDPC label-free images; Section 5: Autofocusing procedure discussion and Figure S3: Schematic of autofocusing procedure; Section 5.1: Duration and Figure S6 - Time required per calculation step; Supporting figure 7: An example of autofocusing curves when the total drift was ∼ 10 *µ*m; Supporting figure 8: QDs tracking with and without active autofocusing in live brain slices.

