## Supplemental information for "Back-illumination Phase Imaging Enables Nanoscale Drift Stabilization in Non-transparent Biological Tissues"

### 1 Comparison of real-time autofocusing methods

Table 1: Comparison of Methods

| Method | Accuracy | Sample types | Duration | Main drawback | Reference |
|---|---|---|---|---|---|
| IR reflection-based PSF | around 20 nm | Thin samples | Limited by sample deformation | Only thin samples fixed on the cover-slip | Nikon Perfect Focus |
| Fiducial-based | < 20 nm | Both thin and thick | Indefinite | Requires presence of fiducial in the plane of interest, which may not always be feasible. Fails for high amplitude drifts | [1, 2, 3] |
| Intensity or DPC images | <6 nm | Thin samples | Limited by sample deformation | Trans-illumination | [4, 5, 6] |
| hDPC-OBM | below 10 nm inside tissue | Thick and/or opaque tissues | Limited by sample deformation | | this study |

### 2 Autofocusing using trans-illumination

We used Quadriwave Lateral Shearing Interferometry (QLSI) camera to generate intensity and quantitative phase images (QPI) of the brain tissue (Figure S1) [9]. QPI images contain the phase information collected along the light path, thus integrating the multiple cell layers with distinct phase profiles. As a result, the QPI provides images containing even less spatial details than intensity (Figure1 (main text), g) leading to flatter CC curve (Figure1 (main text), f, green curve). Moreover, QPI is also affected by uncertainties related to the offset value, which is obtained by integrating signals from interferograms taking into account a reference phase. This manipulation is more difficult to implement in thick live tissues.

Interestingly, it is well known, that phase contrast, and in particular differential phase contrast microscopy (DPC) [10], enables the acquisition of images of biological samples with high degrees of details [11]. Therefore, we hypothesize that images obtained from derivatives of QPI, notably hDPC will be more effective than QPI itself or intensity-based methods.

In practice, we define here hDPC from the two intermediate and orthogonal DPC images along the x and y- axes ($\nabla_x$W and $\nabla_y$W) obtained from interferograms generated by QLSI [9]. The two images are simply subtracted to generate the final hDPC image (FigureS1):

$$hDPC = \nabla_x W - \nabla_y W \qquad (1)$$

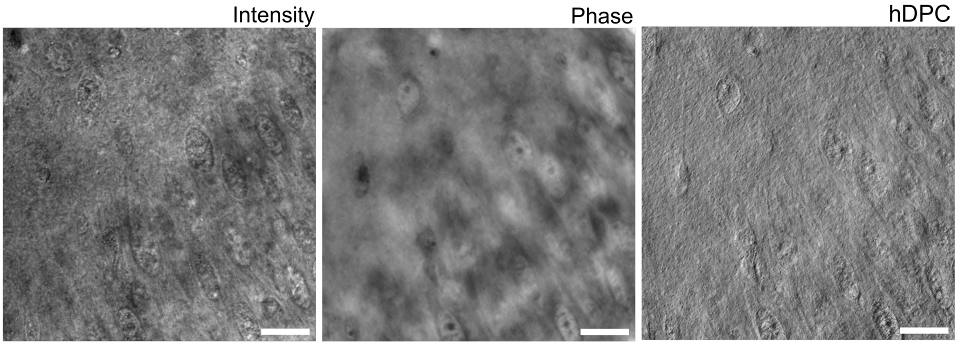


Figure S1: An example of intensity, phase and hDPC images obtained using QLSI module on comercial trans-illumination setup

In Figure S2, we present the bead super-localization signal recorded simultaneously with active autofocusing (FigureS2, c, green curve) and without such stabilization (Figure S2, c, orange curve). As mentioned above, during the autofocusing procedure drift values are calculated right before their correction (Figure S5, b, red dots). Additionally, the position of z-drive of the microscope is monitored (Figure S2, b, blue dots). In these experiments, it was possible to maintain the focus position within a range of ±25 nm over ∼ 8 minutes. These values are at the resolution limit of the z-drive accuracy of the microscope (25 nm). The overall axial drift corrected during the acquisition was 425 nm. Moreover, SD value of the mean axial position (calculated using rolling window over 100 frames) of the fluorescent bead was 11 nm (Figure S2, c, green curve). In comparison, with the autofocusing system deactivated, bead axial position varied by several hundreds of nanometers along the z-axis during the acquisition time due to drifts of the sample (Figure S2, c, yellow curve).

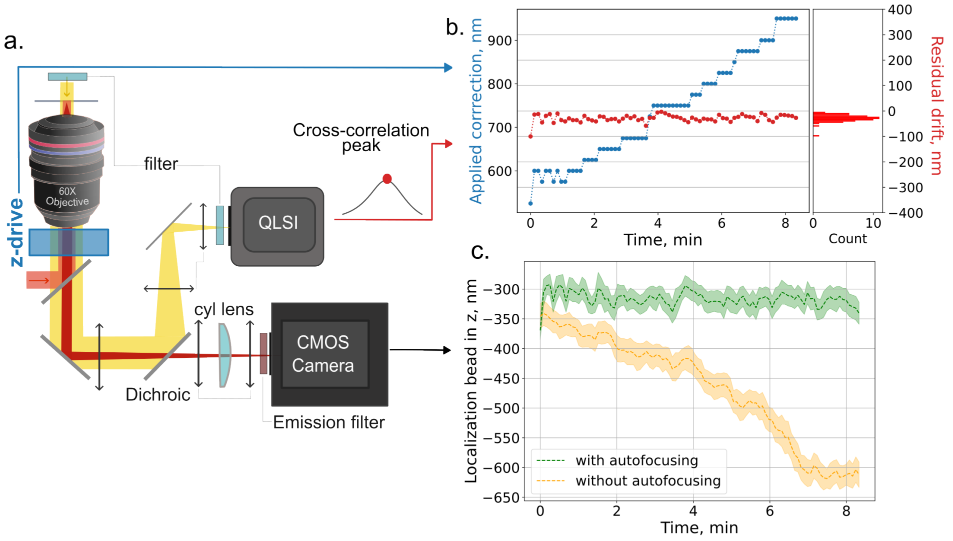


Figure S2: **a.** Schematic of optical setup using transmission illumination. In this schematic, the yellow beam represents white light Köhler trans-illumination, which traverses the filter before illuminating the sample. The light red beam represents laser illumination, while the dark red beam represents the emission detected by the fluorescence channel camera. **b.** During autofocusing the drift was calculated for every $5^{th}$ frame acquired using cross-correlation of gradients images (red curve, axial drift obtained right before its correction). Additionally, the current z-drive position of the microscope was recorded (blue curve). **c.** The bead axial position recorded with and without active stabilization.

**Optical setup for transillumination**

The measurements were performed on commercial inverted microscope Nikon TiE equipped with a 60X 1.3 NA silicon oil immersion objective (Olympus). The sample was illuminated using Köhler trans-illumination with white light from halogen lamp (Nikon) filtered by a bandpass filter (FESH600, Thorlabs) right before the sample. The detection path consisted of two cameras (see Figure S5,a). The white light was separated from the fluorescent signal using a dichroic mirror (FF655-Di01). Fluorescent signal was detected by sCMOS camera (KURO®, Princeton Instruments) controlled with Micro-Manager software [12]. The transmitted white light was detected by a commercial QLSI module (SID4 sC8, Phasics). An additional bandpass filter (FESH600, Thorlabs) was placed right before the QLSI module.

For imaging Tetraspec beads and QDs, same elements as for back-illumination setup describe in main article text were used. The control of white light illumination, z-drive of the microscope and QLSI camera was performed using custom Python based.

In the case of QLSI, a reference image needs to be recorded ones for every sample. Typically, the reference is obtained by moving fast around the sample using longer exposure time ($\sim$ 80 ms). For all the other acquisitions the exposure time was typically chosen to be 10 ms, as introduced by Baffou et al. [9]

#### 3 Features sensitivity to defocus

The detected features depend exclusively on the intrinsic structural content of the tissue within each focal plane. Because tissue architecture varies across tissue types and anatomical regions the distribution and characteristics of the identified features are inherently sample-dependent.

Additionally, to demonstrate the sensitivity of image features to defocus, we introduce a quantitative metric based on the number of matching points between frames within the recorded z-stack, as shown in FigureS3. These graphs were obtained similarly to comparison CC curves (Figure 1f, main text), the number of matching points was calculated for each image in the reference z-stack against the middle image. Accordingly to CC curves, these results confirm that the identified features are highly sensitive to defocus, particularly in hDPC images.

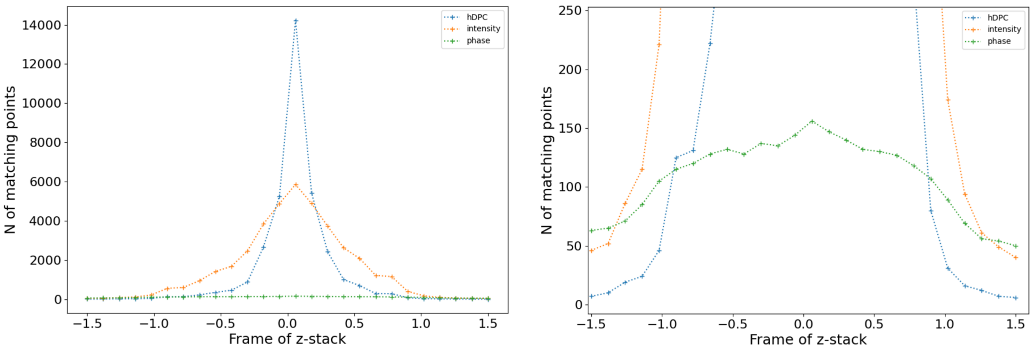


Figure S3: **Number of matching features**

#### 4 xy-drift correction

Lateral drift was corrected numerically at each autofocusing step to prevent additional movement of the slice in the imaging medium and avoid backlash errors from stage adjustments. Additionally, in cases where the speed of axial drift correction is critical, this approach eliminates the need for the waiting time required for mechanical movement of the stage. xy-drift was corrected using SIFT method in Python.

In this study we selected the SIFT method due to its ability to detect scale- and rotation-invariant local features. This enables reliable feature and matching across variations in scale, orientation, and illumination. This method proved its robustness and high performance for noisy images and is freely available in Fiji and Python. A limitation of this method is its high computational cost, which is one of the primary factors slowing down our algorithm.

Interestingly, alternative feature-detection methods could also be applied for drift correction tasks such as SURF (Speeded-Up Robust Features)[7] or ORB (Oriented FAST and Rotated BRIEF) [8]. These methods are less robust than SIFT, but they offer significantly improved computational efficiency and may be worth evaluating in future work. Nevertheless, in the present study, we opted for SIFT because robustness was prioritized over computational speed. A processing time of 1.5-2 seconds per step was considered acceptable for our purposes.

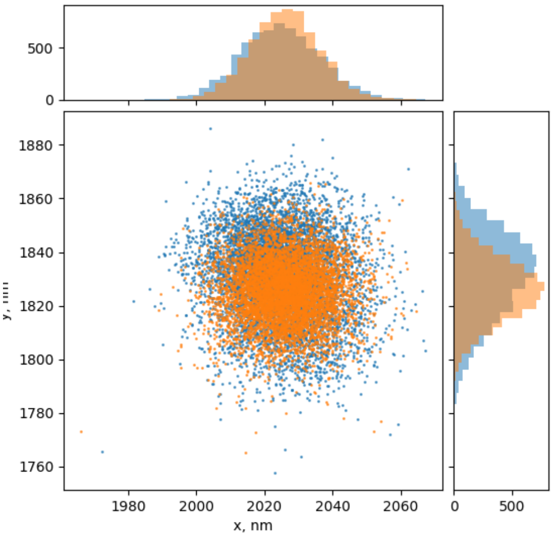


Figure S4: **Correction of xy-drift** of the Tetraspec bead using ThunderSTOMR plugin (orange) (SD_$x$ = 10 nm, SD_$y$ = 12 nm) and SIFT method applied to hDPC (blue)(SD_$x$ = 11 nm, SD_$y$ = 16 nm). In general ThunderSTORM drift correction which uses cross-correlation method performs better on immobile particles or when there is very high number of localizations detected on static structures. However it can not be applied for single particle tracking data.

#### 5 Autofocusing procedure discussion

During the acquisition with active autofocusing it is required to visually examine the shape of CC curves, which is enabled in Python-based software. When distortions in the curve become significant, for example, when the peak becomes highly asymmetric or noisy, this may lead to strong outlier values in the estimated axial position (derived from the CCC peak, which is also displayed in real time). When distortion becomes significant - for example, when the peak becomes highly asymmetric or noisy (see schematic example FigureS5,b) it is necessary to update reference z-stack.

The validity period of a reference z-stack can vary depending on the specific tissue type, region, and experimental conditions. For example, in live brain slices, we limited acquisitions to approximately 10 minutes. Beyond this time, acquisitions are often stopped due to photobleaching of fluorescent particles or rapid morphological changes in some brain regions, making further imaging impractical. Nevertheless, the validity of the reference z-stack in fixed immobile tissues can be much longer.

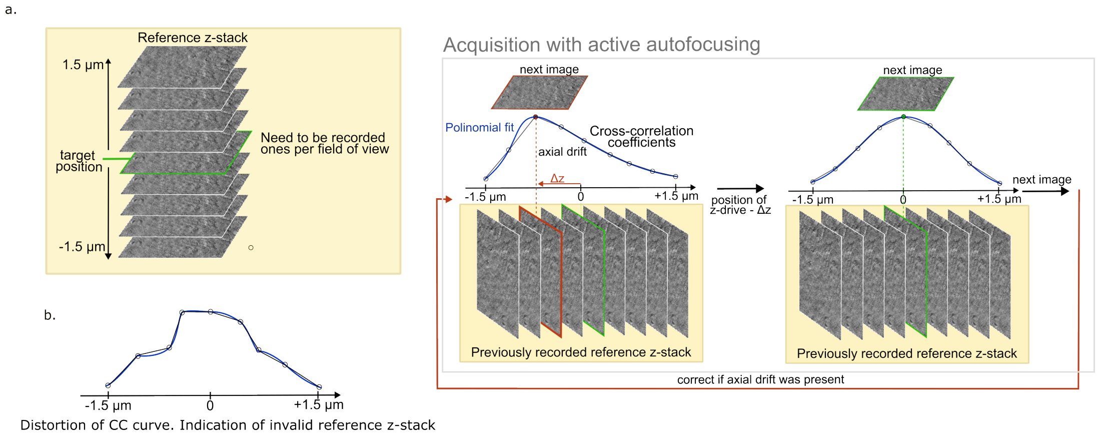


Figure S5: **a.**Schematic of the autofocusing procedure. **b** Schematic example of a distorted CC curve. This indicates that current reference z-stack became invalid.

Additionally, we can estimate velocity and amplitude for specific parameters used in this study:
- time required for each autofocusing step 1.5 s;
- z-stack acquired in range -1.5:1.5 $\mu$m;

Thus, the maximal speed of the sample movement that can be corrected would be 1.5 $\mu$m / 1.5 s = 1 $\mu$m/s.

If the peak of the cross-correlation curve falls outside the correction range ($\pm$1.5 microns), its position is assigned to the nearest extreme value (e.g., +1.5 or -1.5 microns). Consequently, if the drift exceeds this range, it may require multiple autofocus correction steps to return the system to the desired 0 position.

#### 5.1 Duration

The reported autofocus interval includes: camera exposure and readout, calculation of hDPC images, keypoints detection and linear image alignment using these keypoints, CCC calculation and z-drive position adjustment. Reference the z-stack recording performed prior to acquisition with autofocusing, which in our case was completed in 12 seconds (to acquire set of 4 images at each z-position in range -1.5 $\mu$m to 1.5 $\mu$m with a step of 50 nm, resulting in 60 images stack). ROI size used for image processing in this study was 700 x 700 pixels.

For the acquisitions performed in this study we applied the autofocusing every five images leading to $\sim$ 1,5 s between corrections. In principle, the correction time could have been twice faster but we noticed that precision of stabilization would not have been improved. The time required for one autofocusing step can be further reduced by optimizing the code and applying GPU-accelerated calculations using the CUDA library in Python.

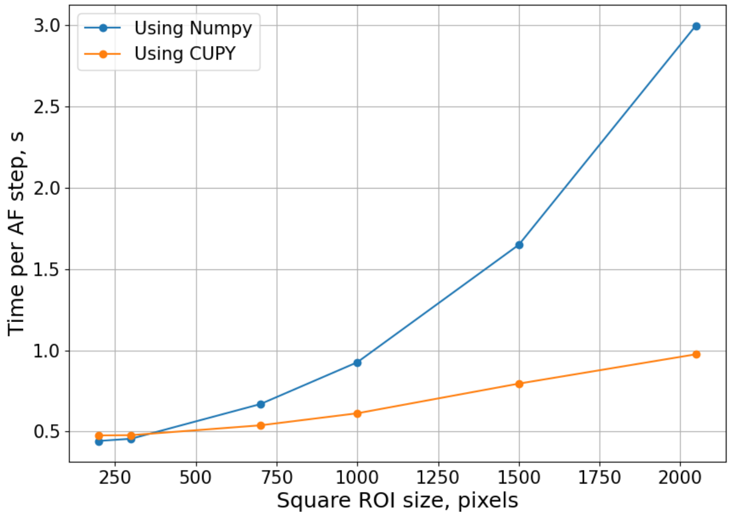


Figure S6: Time required to perform one step pf autofocusing using **a.** Numpy or **b** Cupy llibraries in Python

### 6 Supplementary figure 7

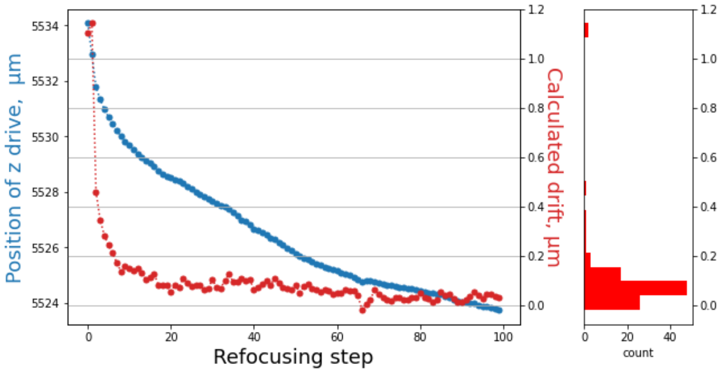


Figure S7: The axial drift calculated right before its correction during the acquisition with autofocusing in live brain slice (red dots). The corresponding position of z-drive of the microscope (blue dots). The total corrected drift was 10 $\mu$m. The SD value of the drift after 10 steps was 34 nm.

### 7 Supplementary figure 8

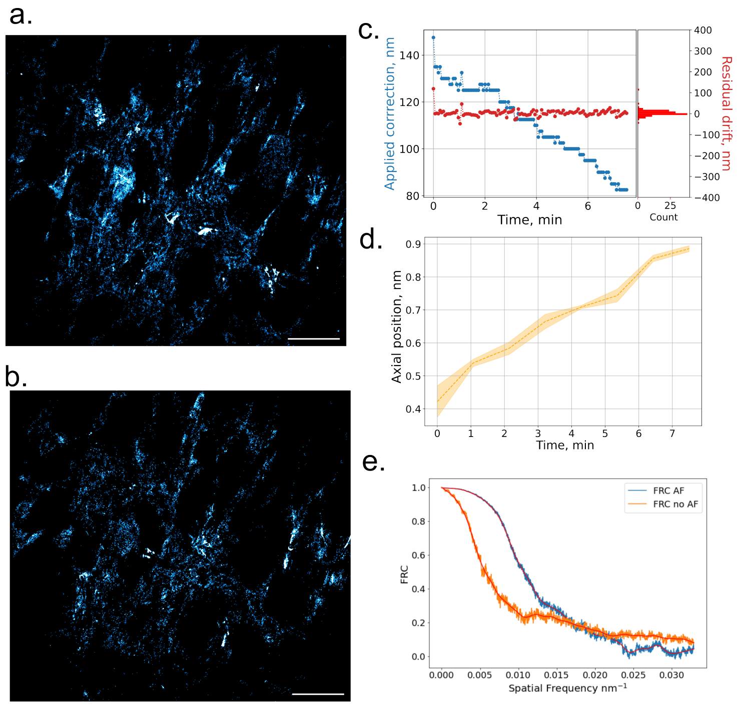


Figure S8: ECS maps generated form QDs localizations with active autofocusing **a** and without **b**.The drift detected (red dots) with SD value of $\sim$ 30 nm and the position read from z-drive (blue dots). The total drift during the acquisition without active autofocusing was ~500 nm **d**. **e** The FRC curves calculated for the images obtained with active autofocusing and without.

[11] Bruce Alberts, Alexander Johnson, Julian Lewis, Martin Raff, Keith Roberts, and Peter Walter. *Molecular Biology of the Cell. Looking at the Structure of Cells in the Microscope.* New York: Garland Science;, 4th edition edition, 2002.

[12] Arthur Edelstein, Nenad Amodaj, Karl Hoover, Ron Vale, and Nico Stuurman. Computer control of microscopes using µmanager. *Curr Protoc Mol Biol*, Chapter 14:Unit14.20, Oct 2010.
